# The extraordinary robustness of mitotic spindle assembly to microtubule dynamics plasticity revealed by end-binding proteins and tubulin tagging

**DOI:** 10.64898/2026.07.29.741283

**Authors:** Morgane L.V. Robert, Aurélien Perrier, Janet Chenevert, Layla El-Mossadeq, Gilliane Maton, Alex McDougall, Anna Castro, Thierry Lorca, Julie C. Canman, Stefania Castagnetti, Julien Dumont, Benjamin Lacroix

## Abstract

Microtubules are dynamic, conserved cytoskeletal filaments that are essential for all eukaryotic cells. Microtubule dynamic properties are primarily characterized by measuring how fast they grow and shrink (growth and shrinkage rates) and how often they switch between assembly and disassembly (catastrophe and rescue frequencies). These four parameters measured for individual filaments can inform on the behavior of the entire microtubule network at the cell level. By comparing microtubule dynamics in *Caenorhabditis elegans* one-cell embryos using different genetically-encoded fluorescent probes, we observed an unexpected high variability in these parameters. Microtubule dynamics parameters were consistently higher in *C. elegans* strains expressing a fluorescently labelled microtubule end-binding protein than in strains relying on tubulin labelling, with microtubule growth rates differing by nearly a factor of two between the two conditions. This discrepancy was not limited to *C. elegans*, as we observed a similar effect in embryos of the tunicate *Phallusia mammillata*. Despite this, spindle size and assembly timing were only mildly affected. However, embryos expressing labelled end-binding protein exhibited higher frequency of mitotic defects and perturbed embryonic development upon exposure to various stresses such as elevated temperature or a compromised spindle assembly checkpoint. Thus, our work reveals both the remarkable robustness of mitotic spindle assembly in response to extreme microtubule dynamics plasticity, and the requirement for strict control of microtubule dynamics across successive early embryonic divisions. Our findings should also serve as a cautionary note when using tagged end-binding proteins to measure microtubule dynamics.

## Introduction

Microtubules are composed of polymers of tubulin heterodimers, which possess a remarkable capacity to self-assemble. The resulting filaments are labile and undergo continuous, stochastic transitions between phases of growth and shrinkage, a behavior known as dynamic instability (Mitchison and Kirschner, 1984). This dynamic property is essential for most microtubule functions in eukaryotic cells, including intracellular transport and chromosome segregation during mitosis. The abrupt transition between growth and shrinkage is called catastrophe, while the inverse phenomenon is named rescue. Measuring frequencies of catastrophe and rescue, as well as the rates of assembly and disassembly, not only reveals the dynamic properties of individual filaments but also provides insights into the size, shape, and behavior of macromolecular networks composed of hundreds to thousands of microtubules, such as the mitotic spindle (Verde et al., 1992). Microtubule dynamics has been measured *in vitro* and *in vivo* in a wide variety of cells, species and biological contexts, and exhibits remarkable variability between species, cell types, and even among microtubules within a single network (Shelden and Wadsworth, 1993; Lacroix et al., 2014; Srayko et al., 2005; Gomes Paim and Bechstedt, 2024). Variation in microtubule dynamics is even observed within a single microtubule during one assembly or disassembly event (Howard and Hyman, 2009; Bollinger et al., 2020; Luchniak et al., 2023; Lacroix et al., 2016). Distinct growth rates within a microtubule subpopulation or during a single growth excursion were noticed in the earliest experiments of microtubule dynamics observation and are in part attributed to intrinsic fluctuations in the assembly configuration of dimers and protofilaments within the microtubule lattice (Howard and Hyman, 2009; Gildersleeve et al., 1992; Chrétien et al., 1995). These variations in microtubule dynamics can be modulated by the composition in tubulin isoforms (Panda et al., 1994; Honda et al., 2017; Pamula et al., 2016; Vemu et al., 2017), by microtubule associated proteins (MAPs) (van der Vaart et al., 2009; Cassimeris, 1993), as well as post-translational modifications of both tubulin and MAPs (Janke and Magiera, 2020). In addition, local environmental factors such as viscosity, crowding (Wieczorek M, 2013; Molines et al., 2022), cytoplasmic density (Geisterfer et al., 2020), spatial variation (Ishihara et al., 2021), temperature (Fygenson et al., 1994; Srayko et al., 2005; Li and Moore, 2020), cell size (Lacroix et al., 2018; Rieckhoff et al., 2020; Geisterfer et al., 2020), or mechanical forces applied on microtubules (Dogterom and Yurke, 1997; Janson et al., 2003) can also modulate microtubule dynamics.

Fluorescent proteins that label either the microtubule lattice (such as tubulin) or the growing ends (such as plus-end binding proteins (EBs)) are widely used to visualize microtubules and measure their dynamic properties in living cells. Tubulin labelling allows measuring the four parameters of microtubule dynamics, whereas the use of EBs, only enables measurement of the polymerization rate and catastrophe frequency (Mimori-Kiyosue et al., 2000; Tirnauer and Bierer, 2000; Schuyler and Pellman, 2001). Although genetically encoded N-terminally or C-terminally tagged tubulins or EBs are the most commonly used microtubule probes *in vivo* (Vicente and Wordeman, 2019; Zwetsloot et al., 2018), chemically labelled purified tubulin can also be injected into cells or embryos of non-model organisms to visualize microtubules and measure their dynamic properties (Chenevert et al., 2024; Wadsworth and Sloboda, 1983). Studies using fluorescently labeled tubulin or EB proteins report microtubule growth rates in 1-cell *C. elegans* embryos ranging from approximately 0.2 µm/s to 1 µm/s, with substantial variability across different studies. This high variability could arise from differences in experimental conditions used to image and quantify microtubule dynamics. However, when comparing these earlier studies, an unexpected trend emerges: average microtubule growth rates fall around 0.4–0.5 µm/s when measured with tagged tubulin (Kozlowski et al., 2007; Labbe et al., 2003; Lacroix et al., 2018), whereas strains expressing tagged EBs report an average growth rate of approximately 0.7 µm/s (Sallee et al., 2018; Srayko et al., 2005; Rodriguez-Garcia et al., 2018). This pattern suggests that the choice of microtubule probe—rather than the experimental setup or measurement method—largely accounts for the observed differences in dynamics, with tagged EBs consistently yielding higher estimates of microtubule growth rate. However, a direct side-by-side comparison of these two microtubule probes under matched imaging conditions and analysis protocols has not yet been performed. Furthermore, although switching from tubulin-to EB-tagged constructs produces striking differences in measured microtubule dynamic parameters, these changes have not been linked to any detectable alterations in animal development or behavior. This leaves unresolved whether the observed variations in dynamics reflect mere measurement artifacts or instead represent functionally relevant changes in microtubule behavior.

In the present work, we documented the difference in microtubule dynamics depending on the strategy used to label and visualize microtubules (EB-or tubulin-tagging). Using eight different worm strains expressing either fluorescently tagged tubulin (tubulin strains), or EB proteins (EB strains), or both, we confirmed that measured microtubule growth rates are higher (by at least 60%) in EB strains than in strains expressing tubulin alone. This discrepancy in measured microtubule growth rate is not specific to *C. elegans* as we observed similar differences in *Phallusia mammillata* ascidian embryos by using micro-injected labelled tubulin or labelled EB. In *C. elegans*, we found that the measured microtubule growth rate was relatively consistent among different tubulin strains independently of the fluorophore used for tagging, the labelled tubulin isotype, or the promoter driving its expression. In contrast, using different fluorescent EBs, expressed ectopically or endogenously tagged by CRISPR/Cas9, we found a higher variability in the average measured microtubule growth rate between conditions. Microtubule catastrophe frequency and shrinkage rate were also higher in these EB strains, resulting in a higher microtubule turnover. All EB strains also presented higher sensitivity of early embryos to thermal stress and exhibited more chromosomal segregation errors during embryonic divisions. On a broader perspective, our work revealed the large amplitude of individual microtubule dynamics that cells can tolerate without exhibiting dramatic defects. As microtubule and spindle assembly are crucial for development and reproduction of metazoan, our work provides additional evidence that flexibility in molecular processes might be crucial to confer robustness at cellular and animal scales (Kirschner and Gerhart, 1998).

## Results

Distinctly tagged microtubule *C. elegans* strains show different microtubule growth rates under identical experimental conditions.

Using spinning-disk confocal live imaging of *C. elegans* 1-cell embryos, we measured microtubule dynamics in eight different strains harboring either single or combinatorial fluorescently tagged tubulins (tubulin strains) or end-binding proteins (EBs, EB strains) (Figure 1 and Table S1). Tubulin strains express only fluorescently tagged *α*-or *β*-tubulin under germline-specific promoters (pie-1 or mex-5), enabling protein expression in oocytes and during early embryonic development (see Table S1). EB strains express CRISPR/Cas9-endogenously tagged or ectopic pie-1 promoter-driven fluorescent end-binding protein (EBP in *C. elegans*), with or without additional tagged tubulin. All strains were imaged under identical experimental conditions (see Experimental Procedures). Since microtubule dynamics vary during the cell cycle (Srayko et al., 2005) and scale with cell size (Lacroix et al., 2018), we focused our measurements on spindle and astral microtubules in the *C. elegans* 1-cell embryo during mitotic spindle assembly i.e., between nuclear envelope breakdown and anaphase onset (Figure 1A and 1B). We first analyzed microtubule growth rates, measurable from microtubule tracks (in tubulin strains) or EB comets (in EB strains) (Figure 1 A-C). In strains expressing both fluorescent markers, the growth rate was measured using the tagged-tubulin marker. In line with previous reports (Srayko et al., 2005; Lacroix et al., 2014, 2018), microtubule growth rates vary widely across strains and even within individual embryos, resulting in large standard deviations; astral microtubule rates exceed those of spindle microtubules. In all strains, microtubule growth rates followed a unimodal log-normal distribution with longer tail toward higher rates, typical of cellular reaction processes (Furusawa et al., 2005). Consistent with prior literature, *C. elegans* strains broadly segregated into two distinct categories based on their average microtubule growth rates. Tubulin strains exhibited average microtubule growth rates of ∼0.3 µm/s for spindle microtubules and ∼0.4 µm/s for astral microtubules (Lacroix et al., 2018; Labbe et al., 2003; Kozlowski et al., 2007). EB strains showed higher microtubule growth rates, averaging 0.49-0.58 µm/s for spindle microtubules and 0.55-0.79 µm/s for astral microtubules (Sallee et al., 2018; Srayko et al., 2005; Rodriguez-Garcia et al., 2018). Importantly, we previously demonstrated that microtubule growth rates during early embryonic development are identical across three strains expressing GFP-or mCherry-tagged *β*-tubulin under distinct promoters (Lacroix et al., 2018). Here, we further show that microtubules exhibit identical assembly dynamics in *C. elegans* strains expressing either GFP-*α*-tubulin or mCherry-*β*-tubulin. These results confirm that microtubule growth rates remain robustly consistent across tubulin strains, regardless of isoform (*α* or *β*), promoter, or fluorescent tag (GFP or mCherry). Interestingly, average growth rates were similarly high in CRISPR/Cas9-engineered EB strains and those with ectopic pie-1 promoter-driven tagged EB. Independent of average growth rates, individual microtubule growth rate distributions exhibited more variability in EB strains, as indicated by larger standard deviations (Figure 1 D and E). One potential explanation for these growth rate differences is that tagged EBs and tubulin reveal distinct subpopulations of microtubules. However, in EB strains co-expressing tagged tubulin, higher rates persisted even when tracking microtubules via the tubulin signal (magenta and cyan dots in Figure 1D, E). Furthermore, velocities measured by simultaneous two-color imaging of both markers were identical (Figure S1), ruling out the possibility that tubulins and EBs label distinct microtubule subpopulations with different dynamics. Our results indicate that tubulin tagging yields more consistent and reliable results across strains. The presence of tagged EBs in *C. elegans* embryos appears to alter microtubule dynamics by increasing microtubule growth rate.

**Figure 1.**
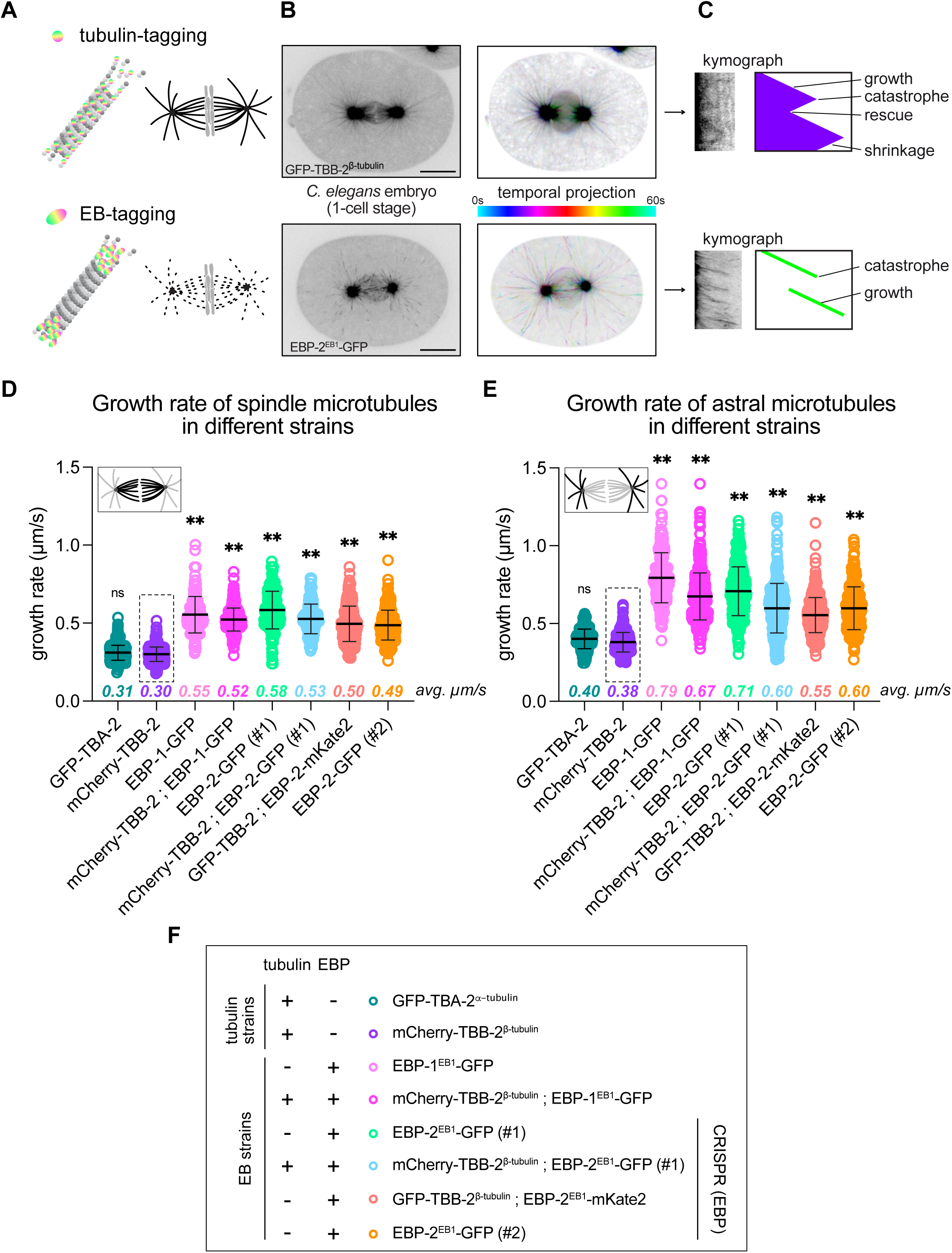
Microtubule growth rate of strains expressing tagged-tubulin or tagged-EBs. (A) Schematic representation of the two main methodologies used to visualize microtubules by fluorescence in *C. elegans* embryos. (Top) Fluorescent-labelling of tubulin which incorporates in the lattice to label the fibers. (Bottom) Fluorescent-labelling of an end-binding protein (EB) decorates the growing tips of microtubules. Centrosomes contain a high number of microtubules plus ends and minus ends. EB binding to growing plus end generates fluorescent “comets”. (B) Left: one image from a timelapse video of a representative *C. elegans* 1-cell embryo from a tubulin-tagged (top) or EB-tagged strain (bottom). Right: temporal projections showing microtubule trajectories. Scale bars correspond to 10 µm. (C) Kymographs extracted from microtubule trajectories to visualize and quantify microtubule dynamics. (D) Spindle microtubule growth rates measured in 1-cell embryos during metaphase in all strains used in this manuscript. Each circle corresponds to an individual growth event. At least 200 microtubules were analyzed in at least 8 different embryos for each strain. Dashed line rectangle indicates the reference tubulin strain used for statistical comparisons. ns: no significant difference, **: P value <0.0001 in Ordinary one-way ANOVA with Dunnett’s multicomparison test. (E) Same as (D) for astral microtubules. (F) Genotypes for tubulin strains and EB strains used in the manuscript.

Differences in measured microtubule growth rates between tagged tubulin and tagged EB reporters are not unique to *C. elegans*.

We next tested whether the differences in microtubule growth rates measured using tubulin or EB-tagging are specific to *C. elegans*. Indeed, unlike most other organisms studied for microtubule dynamics, *C. elegans* exhibits fast growth rates but short microtubule lifetimes (Chaaban et al., 2018). The microtubule growth rate reported for *C. elegans* 1-cell embryos (>0.6 µm/s) is the highest measured across species (Chaaban et al., 2018; Lacroix et al., 2014; Gigant et al., 2017). This rate even exceeds the theoretical maximum predicted by the diffusion-limited growth (Odde, 1997). This’record’ rate stems primarily from *C. elegans* microtubules’ reduced protofilament number (11 vs. 13 in most species, (Chaaban and Brouhard, 2017) and the high free energy of tubulin dimers promoting assembly (Chaaban et al., 2018). Thus, the discrepancy we observed between tubulin and EB-based microtubule growth rate measurements may arise from these *C. elegans*-specific features. To test this, we measured the microtubule growth rate during embryonic development in a Chordata species, the ascidian *Phallusia mammillata* (Figure 2A). Oocytes were microinjected pre-fertilization with mRNAs encoding EB3-GFP or venus-tubulin, or with chemically labeled porcine tubulin, as previously described (Chenevert et al., 2024). We then measured microtubule dynamics from confocal microscopy movies of 8-cell stage blastomeres. Since *P. mammillata* embryos are larger than *C. elegans*, their 8-cell stage blastomeres have volumes comparable to a *C. elegans* 1-cell embryo. As in *C. elegans*, *P. mammillata* embryos expressing EB3-GFP displayed a higher microtubule growth rate (0.48 µm/s) than those expressing venus-tubulin (0.40 µm/s, Figure 2B), suggesting this tubulin-EB discrepancy is broadly conserved across metazoans.

**Figure 2.**
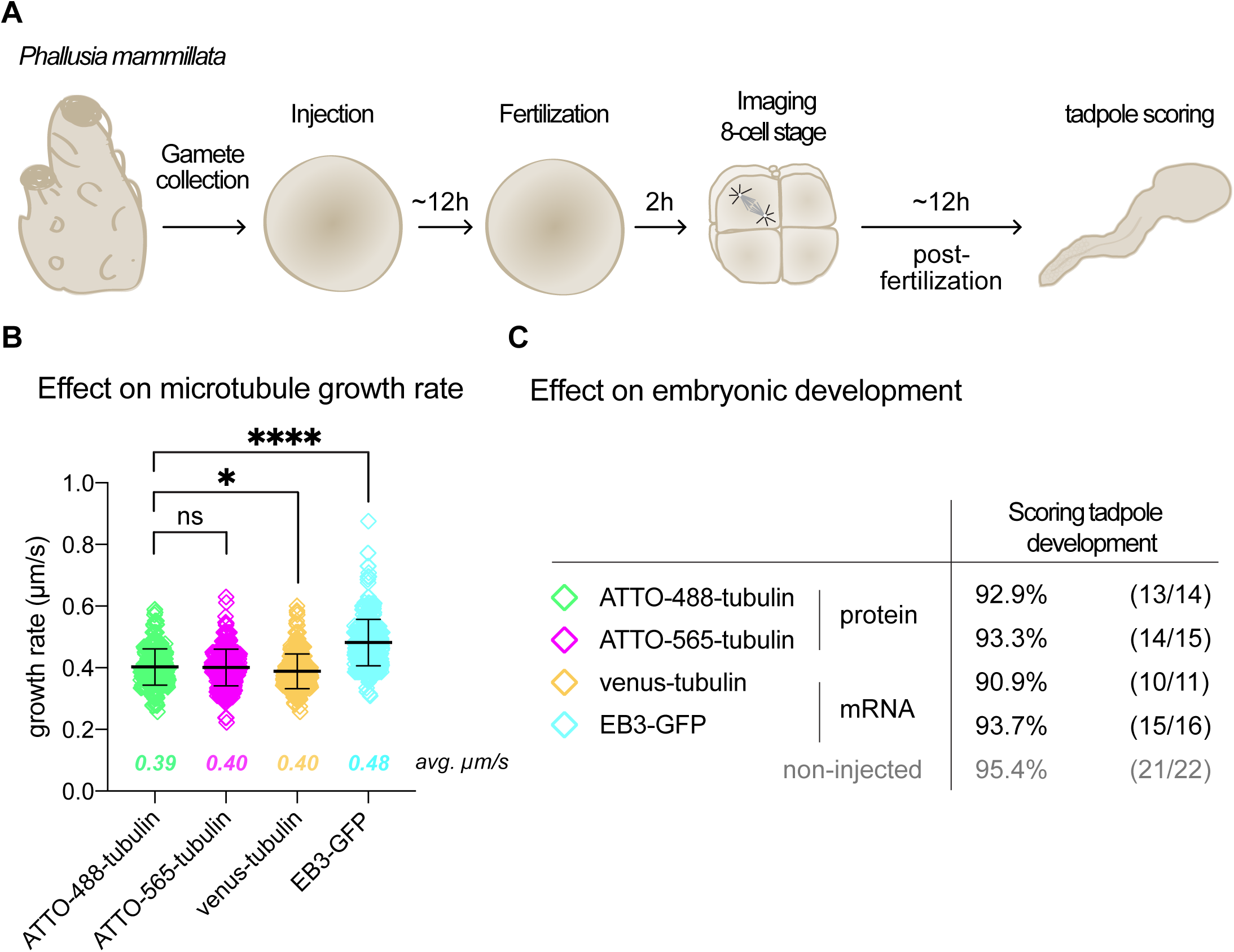
Difference in microtubule dynamics between tubulin-or EB-labelling is conserved in *Phallusia mammillata*. (A) Schematic of experimental procedures for *P. mammillata* embryos. (B) Average microtubule growth rate from at least 9 different embryos and >370 microtubule tracks. Tukey’s multicomparison test was performed. **** P value <0.0001, * P value <0.05. Microtubule growth rate in embryos expressing venus-tubulin was slightly slower (-0.014 µm/s) than the average growth rate measured in embryos injected with ATTO-labelled tubulin. Average microtubule growth rate in embryos expressing EB3-GFP is higher (+ 0.08 µm/s) compared to ATTO-488-tubulin injected animals. (C) Percentage of embryos reaching the tadpole after mRNA or labelled-tubulin injection followed by live imaging. Scoring was performed 12h at 18°C post imaging. Number of embryos scored is given in parentheses for each condition.

Tagged tubulin and EBs induce distinct microtubule turnover rates without altering average microtubule length during spindle assembly.

To further dissect microtubule dynamics differences between tubulin and EB strains, we analyzed all four spindle microtubule dynamics parameters (growth rate, shrinkage rate, catastrophe frequency and rescue frequency) during the first five cleavages of *C. elegans* embryonic development. We selected one representative strain from each category (tubulin and EB strains), both expressing mCherry-*β*-tubulin under the pie-1 promoter (Table S1). The EB strain also expresses GFP-EBP-1^EB1^ under the same promoter. Both strains were outcrossed to wildtype N2 worms to minimize genetic background differences while retaining fluorescent markers, maintained under identical conditions, and imaged using the tubulin signal (mCherry-tubulin, Figure 3A). Microtubule dynamics analysis revealed that, beyond microtubule growth rate differences, both catastrophe frequency and shrinkage rate were elevated in the EB strain (Figure 3C). Rescue frequency remained comparable between strains, except at the 4-cell stage. Overall, microtubule turnover was thus higher in the EB strain than in the tubulin strain, as shown by the larger area of its diamond graphs representing all four microtubule dynamics parameters (Figure 3D). Using the mathematical model of Verde et al. (Verde et al., 1992) and the dynamic parameters of individual microtubules under each condition, we determined the average population growth rate (*J*) for each given microtubule subpopulation (Figure S2A), which corresponds to the rate at which an entire microtubule population grow. This analysis confirmed that microtubules in both strains remained in the expected “bounded” regime typical of mitotic cells, where average length is constrained primarily by high catastrophe frequency (Verde et al., 1992). In this regime, the average length (*L*) of the microtubule population can be estimated from individual dynamics parameters (see equation in Figure 3E, (Verde et al., 1992)). Surprisingly, predicted average microtubule lengths were nearly identical between the tubulin and EB strains despite their markedly different growth rates and turnover (Figure 3E and 3F). These results highlight the remarkable plasticity of microtubule length regulation via dynamics modulation during *C. elegans* embryonic development.

**Figure 3.**
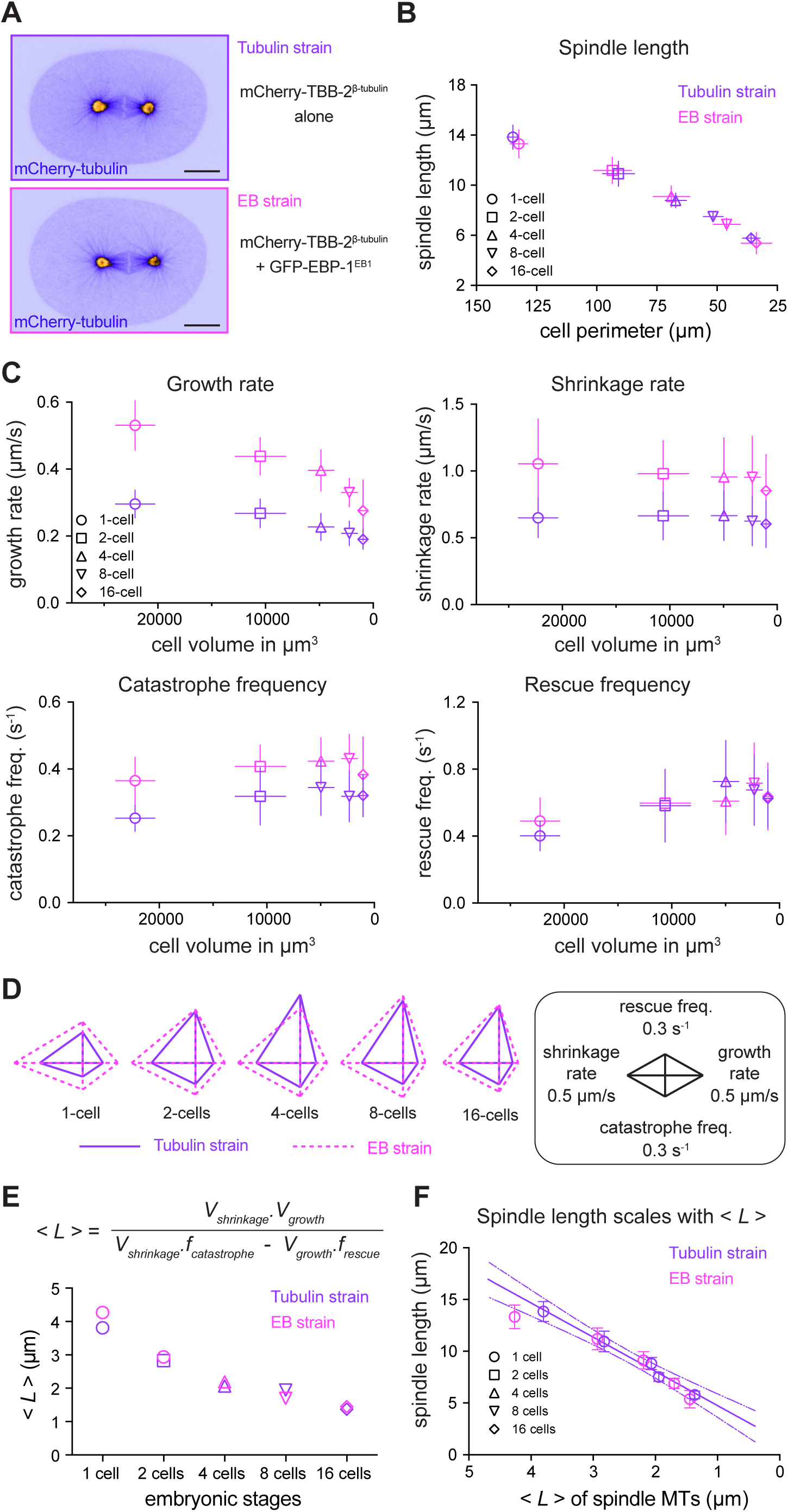
Microtubule dynamics in EB strains are increased compared to tubulin strains with no apparent effect on microtubule lengths. In the entire figure purple corresponds to the tubulin strain and magenta to the EB strain. These strains were also represented in Figure 1 with the same color code. (A) One image from time lapse video in the tubulin channel (mCherry-tubulin) in the tubulin strain (top) and the EB strain (bottom). Scale bar = 10 µm. (B) The graph shows spindle length measured at anaphase onset in both strains and over the 5 first embryonic cleavages. (C) All four microtubule dynamics parameters for spindle microtubules in both strains from the 1-cell to 16-cell stage. Error bars represent standard deviations. Parameters are represented relative to the average cell volume extracted in our previous study (Lacroix et al., 2018). (D) Diamond graphs represent the four parameters of microtubule dynamics in the two strains. Parameters with the same unit are jointly normalized. (E) The average length of spindle microtubules predicted from dynamic parameters following the mathematical model provided by Verde et al. (Verde et al., 1992), given that during the mitotic phase microtubules remain in the bounded regime. (F) Spindle length as a function of the average microtubule length <*L*> estimated in (E). Error bars correspond to standard deviation. Lines represent a linear regression (plain purple) of tubulin strain values with the 95% confidence interval (dashed lines). R squared: 0.9916.

### Microtubule dynamics in tubulin vs EB strains differentially respond to MAP depletions

Given the marked differences in individual microtubule dynamics between the tubulin-and EB-labeled strains, we examined whether the activity of microtubule-associated proteins controlling microtubule assembly and disassembly also depended on the labeling strategy (Jonasson et al., 2020). We therefore depleted three conserved mitotic MAPs, KLP-7^MCAK^, ZYG-9^TOGp^, and CLIP-1^Clip170^, in both strains and assessed their effects on individual and collective microtubule behaviors. These three MAPs control microtubule during mitosis in a wide range of species and cell types (van der Vaart et al., 2009; Bowne-Anderson et al., 2015; Goodson and Jonasson, 2018; Al-Bassam and Chang, 2011). Consistent with previous reports (Srayko et al., 2005), KLP-7^MCAK^ and ZYG-9^TOGp^ depletion slowed microtubule growth (Figure 4A) and shortened spindle length (Figure 4B) in both strains. KLP-7^MCAK^ depletion reduced microtubule growth rates similarly across strains relative to controls (-35.5% in tubulin;-37.6% in EB). (Figure 4A). Surprisingly, ZYG-9^TOGp^ depletion decreased growth proportionally more in the EB (from 0.44 to 0.18 µm/s; ∼59%) than tubulin strain (0.31 to 0.19 µm/s; ∼39%), yielding no significant post-depletion difference. CLIP-1^Clip170^ depletion was previously shown to boost microtubule growth rate in *C. elegans* 1-cell embryos (Rodriguez-Garcia et al., 2018). It had no significant effect on mitotic spindle length in either strain (Figure 4B). Strikingly, CLIP-1^Clip170^ depletion had opposite effects on spindle microtubule growth rates in tubulin and EB strains. Microtubule growth rate increased +37.5% in the tubulin strain but decreased-4.7% in the EB strain. We previously showed that MAP depletion can oppositely affect microtubule dynamics within the same *C. elegans* cell lineage across developmental stages (Lacroix et al., 2014). Here, CLIP-1^CLIP170^ depletion exerts opposite effects in the same 1-cell embryo depending solely on microtubule labeling strategy. Notably, these divergent growth rate changes (+37.5% tubulin;-4.7% EB) converge on identical spindle lengths, revealing deep compensatory mechanisms. Overall, these results show that MAP depletion can exert distinct, and even opposite, effects on microtubule dynamics depending on the initial growth rates and/or the labelling strategy used.

**Figure 4.**
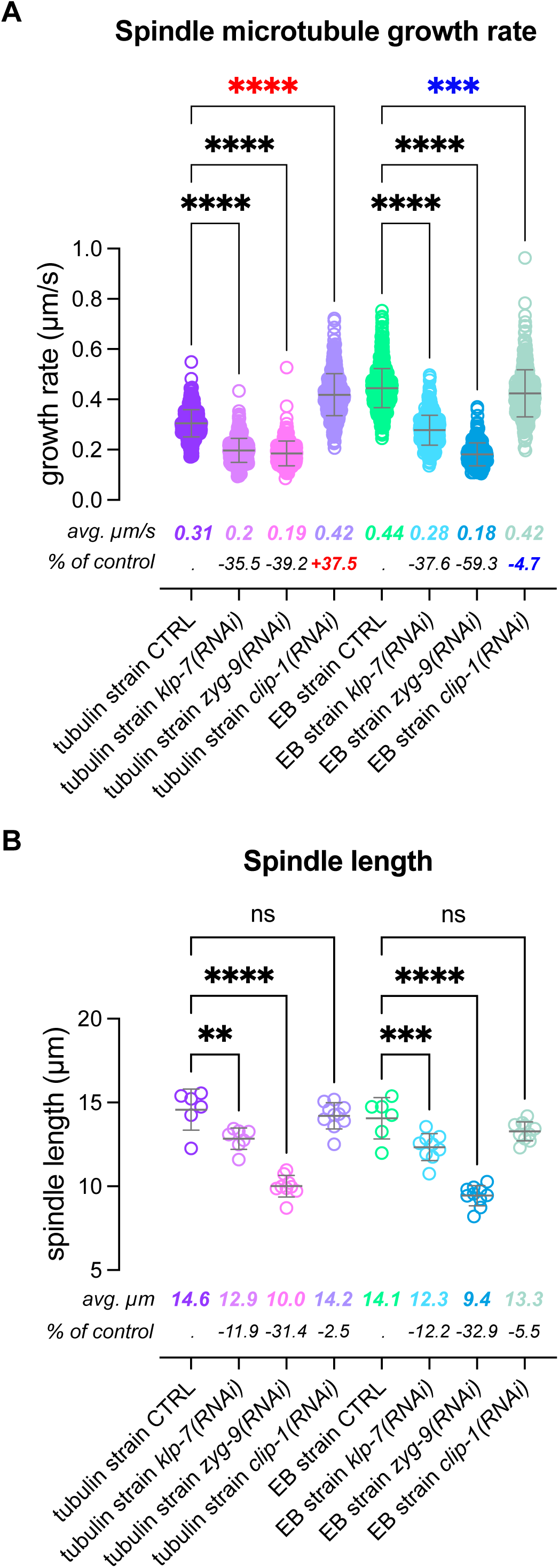
Strain dependent effect of microtubule associated protein depletion. (A) Spindle microtubules growth rate was measured in 1-cell embryo during spindle assembly upon depletion of canonical microtubule-associated proteins: KLP-7^MCAK^, ZYG-9^TOGp^, and CLIP-1^Clip170^. Tubulin strain corresponds to worms expressing mCherry-TBB-2 under embryonic promoter and EB strain contains in addition a GFP inserted in N-terminal of the EBP-2 endogenous locus (by CRISPR/Cas9 editing). Both strains were imaged in the mCherry channel (tubulin). The average growth rate is indicated in µm/s in the respective color. Percentage of change (increase or decrease) relative to the untreated control worms are indicated below. Each circle corresponds to a single growth excursion. Bars represent mean and standard deviation from at least 282 measurements in at least 8 different embryos per condition. Brown-Forsythe and Welch ANOVA test with Dunnett’s multiple comparison. ***: P value < 0.01, ****: P value < 0.001. clip-1(RNAi) induces a significant increase (red****) or significant decrease (blue***) in a tubulin strain (mCherry-TBB-2^tubulin^) and an EB strain (mCherry-TBB-2^tubulin^; GFP-EBP-2^EB1^) respectively. Microtubule growth rate did not differ between tubulin and EB strain upon ZYG-9 depletion (P value = 0.997). (B) Spindle length was assessed in the same condition as (A). Each circle corresponds to the spindle length in a single 1-cell embryo. Bars represent mean and standard deviation. Ordinary one-way ANOVA with Dunnett’s multiple comparison test was performed. ns: no significant difference, **: P value < 0.01, ***: P value < 0.001, ****: P value < 0.0001.

### EB-tagging alters spindle size and assembly timing in the *C. elegans* 1-cell embryo

We next assessed phenotypic traits potentially affected by altered microtubule dynamics. Using movies from growth rate analysis (Figure 1), we measured spindle length (distance between centrosomes) and assembly duration, defined here as the interval from nuclear envelope breakdown (NEBD) to anaphase onset (Figure 5A and (Lacroix et al., 2018). In the reference tubulin strain (mCherry-tubulin), average spindle length was 13.88 µm and was slightly but significantly reduced in most EB strains, except those with endogenously mKate2-tagged EBP-2^EB1^ or co-expressing mCherry-tubulin with ectopically expressed EBP-1^EB1^-GFP (Figure 5B). Spindle assembly duration was significantly shorter in all EB strains compared to tubulin strains. We used differential interference contrast (DIC) microscopy to track centrosomes and nuclear envelope dynamics, comparing spindle size and assembly timing in tubulin and EB strains with the unlabeled wild-type reference (Bristol N2), from which both derive ((Farhadifar et al., 2015) and Figure 5 and Figure S3, see Table S1). We confirmed that both spindle length and assembly duration were similar in the N2 wild-type strain and tubulin strain, but were significantly reduced in the EB strain (Figure S3). Thus, tagged EB expression in the *C. elegans* 1-cell embryo shortens spindle length and accelerates spindle assembly.

**Figure 5.**
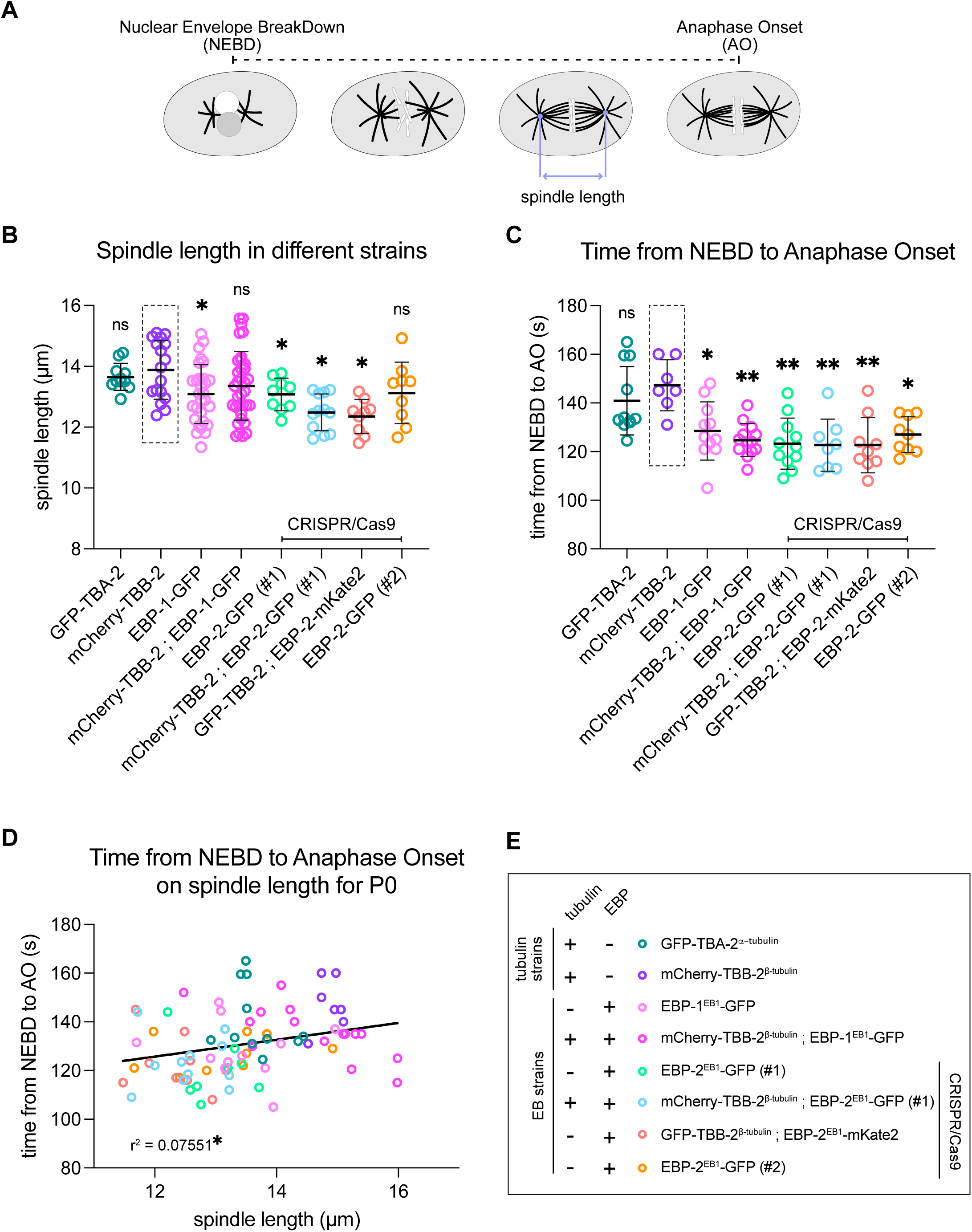
Tubulin strains and EB strains exhibit different spindle length and assembly durations. (A) Spindle assembly duration and spindle length were evaluated for all strains used in this worm from fluorescence confocal live imaging. Spindle assembly duration was defined as the time from nuclear envelope breakdown (NEBD) to anaphase onset (AO). (B) Spindle length in 1-cell embryo measured in all strains used in this study. Dots represent individual embryos. Bars correspond to average and standard deviation of at least 7 embryos per condition. (C) Spindle assembly duration reported for all 1-cell embryos imaged in (B). Bars correspond to average and standard deviation of at least 7 embryos per condition. (B and C) Ordinary one-way ANOVA with Dunnett’s multiple comparison test was performed. ns: no significant difference, *: P value < 0.05, ***: P value < 0.01. (D) Correlation between spindle assembly duration measured in (C) and spindle length (B) for each individual embryo. Every circle corresponds to a single 1-cell embryo in which both spindle length and spindle assembly duration was measured. Line corresponds to a simple linear regression fit, R squared: 0.075. Pearson correlation indicate a significant correlation with a correlation coefficient r = 0.27 and with a P value of 0.013. (E) Legend and color code use in (B, C and D). + and – indicate the presence or absence of the marker (labelled-tubulin or labelled-EB) in all strains.

### *C. elegans* EB strains exhibit elevated thermosensitivity

Having observed cellular effects of EB versus tubulin tagging, we next tested whether these labelling strategies impact embryonic development or animal physiology. Despite the different microtubule dynamics, spindle length, and assembly duration between *C. elegans* tubulin and EB strains, we detected no discernible behavioral differences under standard laboratory conditions (23°C). Similarly, comparable proportions of *P. mammillata* embryos reached the tadpole stage after tagged-EB or tagged-tubulin expression/injection (Figure 2A and C). To unmask potential phenotypes, we varied culture temperatures from 13 °C to 26 °C, a physiologically viable range for *C. elegans*, and quantified embryonic lethality (Figure 6A). As expected, wild-type N2 showed modest temperature-induced lethality increases (Figure 6B, dark grey). Tubulin strain (ectopically expressing mCherry-tubulin) showed a further elevation, but all EB strains, whether ectopically expressing or endogenously tagged, exhibited markedly higher embryonic lethality above 23°C versus wildtype, reaching up to 80% in the strain ectopically expressing an extra copy of EBP-1^EB1^ alongside tagged tubulin. These findings indicate that tagged EBs, or their associated higher microtubule turnover, heighten embryonic thermosensitivity.

**Figure 6.**
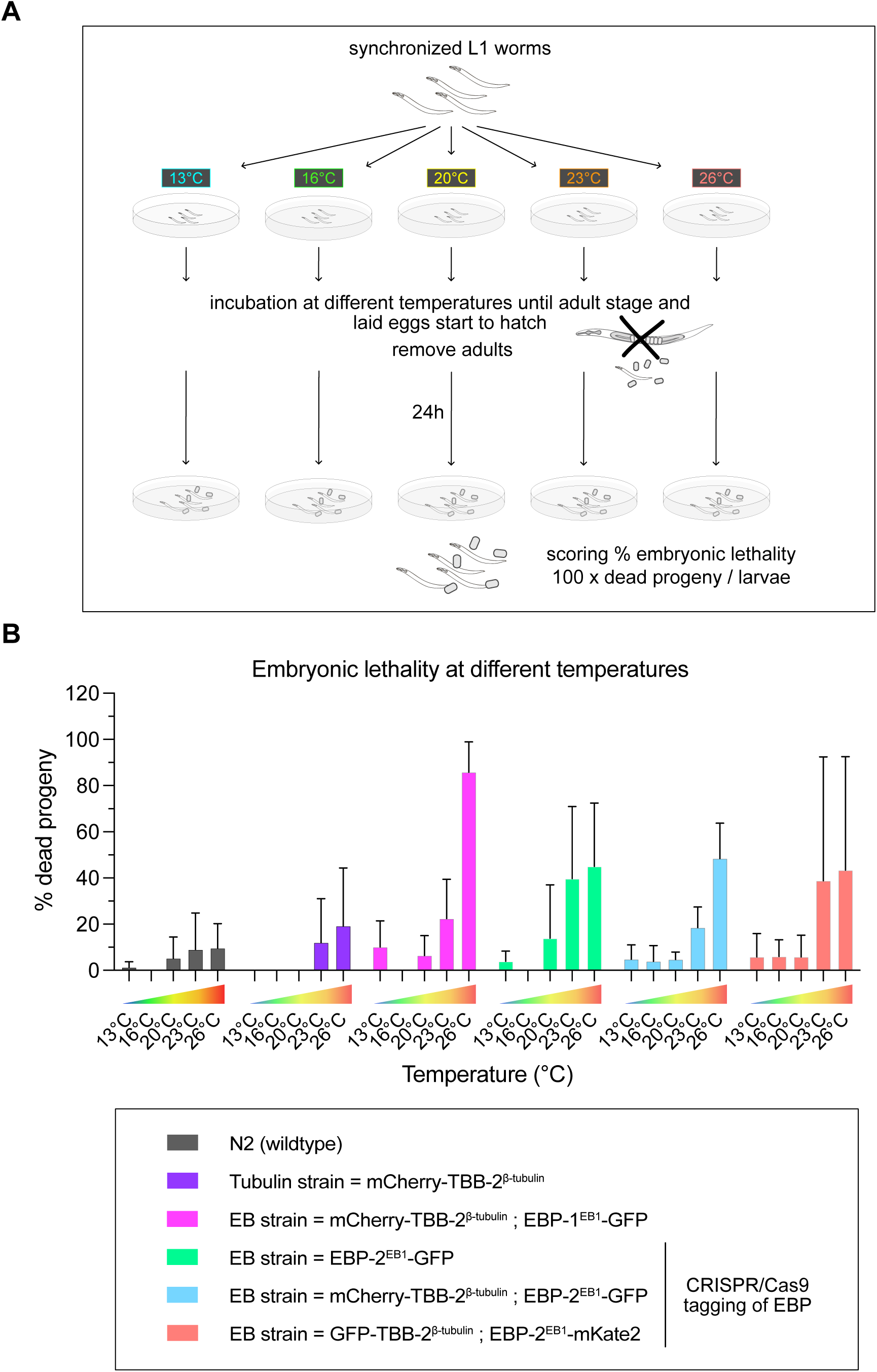
EB strains display increased thermosensitivity. (A) Illustration of the experimental procedure to assess the effect of temperature on embryonic viability of worm strains. (B) Percentage of dead progeny from worms reared at different temperatures from their L1 larval stage. Histograms represent the average of 10 biological replicates per condition, error bars correspond to standard deviation. Bar colors correspond to the different strains indicated in the legend.

### *C. elegans* embryos expressing tagged EB are more prone to mitotic defects

Increased embryonic lethality in EB strains at higher temperature may arise from chromosomal abnormalities (Kyogoku, 2025; Lee and Kiessling, 2017; Zipperlen et al., 2001). Indeed, elevated microtubule turnover has been linked to chromosomal instability in mammalian cells (Ertych et al., 2014). Combined with the shortened mitosis duration observed in EB strains (Figure 5 and S3), this could predispose cells to chromosome segregation errors (Wollman et al., 2005). To assess chromosome alignment and segregation fidelity, we imaged embryos expressing a fluorescent H2B histone marker (mCherry-HIS-58^H2B^) alone or with tagged tubulin/EB, and we quantified alignment defects and lagging anaphase chromosomes at different stages of development (Figure 7A and B). To minimize phototoxicity, we dissected gravid hermaphrodites to obtain embryos at various developmental stages and imaged only single mitoses (<10 min). Under these optimal imaging conditions, chromosome segregation defects were infrequent but detectable. Notably, the frequency of segregation defects increased across all strains during later embryonic divisions (from the 4^th^ cleavage/8-cell stage onward), yet was significantly higher and emerged earlier in the EB strain compared to the tubulin or H2B-only strains (Figure 7C). Thus, the elevated microtubule turnover, or reduced spindle assembly duration, or a combination of both defects appears to increase the incidence of chromosomal abnormalities, potentially explaining reduced embryonic viability in EB strains.

**Figure 7.**
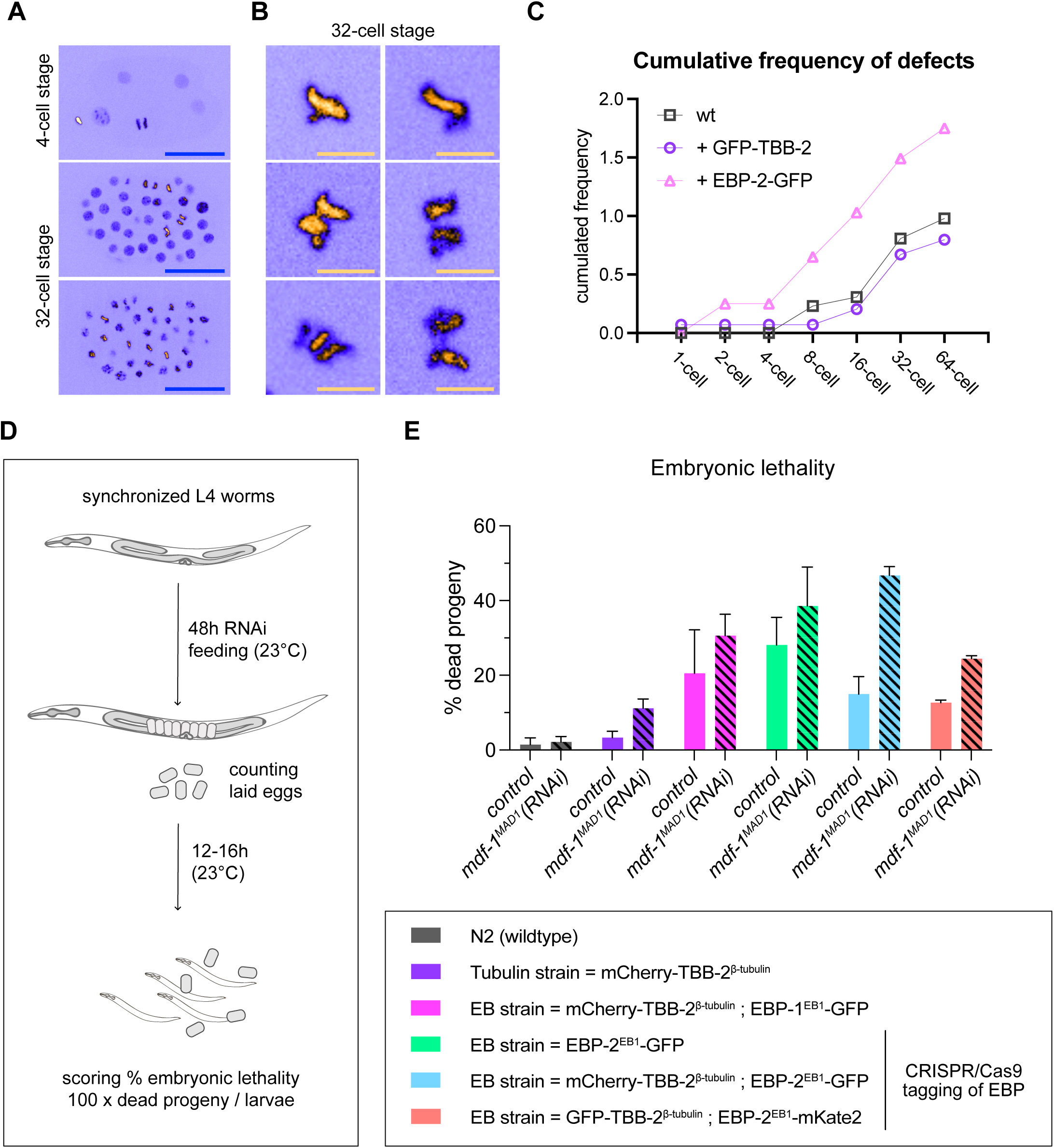
Assessing chromosome segregation errors and SAC sensitivity in tubulin strains and EB strains. (A) Images represent maximal intensity projection of 4 to 6 focal plane of time point from a 3D time lapse video of fluorescent histones (mCherry-HIS-58^H2B^) in *C. elegans* embryos. Top panel corresponds to a 4-cell stage embryo and bottom panels to 32-cell stage. Blue scale bar = 20 µm. Fluorescent signal intensity is represented using the ICA look-up table from Fiji (B) Cropped images from 32-cell stage imaging focused on chromosome alignment or segregation defects: misaligned on the two top images and missegregation of aberrant anaphase features on the bottom panels. Yellow scale bar = 3 µm. (C) Graph of the cumulative frequency of lagging chromosomes in the “wildtype” strain containing only the tagged histone, tubulin strain (GFP-TBB-2^tubulin^) and EB strain (EBP-2^EB1^-GFP). (C) Experimental design to assess embryonic lethality upon depletion of the spindle assembly checkpoint protein (MDF-1^MAD1^). (D) Embryonic lethality at 23°C upon MDF-1^MAD1^ depletion in different worm strains. Colors correspond to the different strain listed in the legend below. Dashed bars highlight dsRNA treatment for MDF-1^MAD1^ depletion. Error bars correspond to standard deviation from 3 independent experimental replicates.

In most cell types, the spindle assembly checkpoint (SAC) prevents chromosome segregation errors by delaying anaphase onset until proper bipolar attachments form. The SAC is weak during early *C. elegans* cleavages (Encalada et al., 2005) but strengthens as cells progressively become smaller across successive divisions (Galli and Morgan, 2016). To exacerbate EB-induced defects, we conducted a synthetic lethality assay by depleting the SAC protein MDF-1^MAD1^ (Figure 7D). In line with previous findings, MDF-1^MAD1^ depletion alone did not increase embryonic lethality in wild-type N2 worms (Figure 7E, (Encalada et al., 2005; Galli and Morgan, 2016)). However, it sharply increased embryonic lethality in all fluorescent strains (tubulin or EB), with similar relative rises despite higher basal embryonic lethality in EB strains. The effect was most pronounced in the dual-labelled strain (Figure 7E, cyan bars). These data suggest that microtubule fluorescent markers, and especially tagged EBs, induce chromosome segregation errors, which the SAC normally mitigates.

## Discussion

### The increased microtubule growth rate and turnover in EB strains affect spindle functions

In this work we investigated how the two most common strategies used to visualize and track microtubules *in vivo* (i.e., tubulin-tagging or tagging of end-binding proteins) impact the measured microtubule dynamics in two distinct species. We showed that the discrepancies in microtubule growth rates reported in the literature were not due to lab specific imaging conditions or quantification bias. Instead, we found that under the same laboratory and imaging conditions, the average growth rate quantified in *C. elegans* and *P. mammillata* embryos using tagged EBs was significantly increased when compared to those measured using tagged tubulin. With the currently available technologies for live imaging in large specimens such as embryos, it is impossible to visualize and measure microtubule dynamics using label-free techniques and therefore to estimate their intrinsic dynamics.

### Altered microtubule turnover and embryonic viability in EB strains

Our results suggest that the increased embryonic lethality observed in EB strains under stress conditions arises from mitotic defects driven by elevated spindle microtubule turnover. We previously reported that individual microtubule growth rate scale with cell size during embryonic development in *C. elegans* (Lacroix et al., 2018). Here we observed that population growth rates (*J,* (Verde et al., 1992)) of both spindle and astral microtubules scale with spindle length from 1-to 16-cell stage (Figure S2B). Interestingly, and as previously reported for individual microtubule growth rates (Lacroix et al., 2018), the population growth rates of astral and spindle microtubules tend to converge to similar values as cells become smaller in tubulin strains. In contrast, this was not observed in EB strains, where astral microtubules remained distinct from spindle microtubules throughout development (Figure S2B). As shown in microtubule dynamics phase diagrams (Figure S2C), EB strains exhibit a shift in dynamic behavior compared to tubulin strains, moving from the limited growth typical of mitosis toward the sustained, “infinite” growth characteristic of interphase (Verde et al., 1992; Shaebani et al., 2016). This effect is particularly pronounced for astral microtubules and is primarily driven by an increased ratio of rescue to catastrophe frequency in EB strains, as cells get smaller, whereas this ratio remains constant in tubulin strains. These altered dynamics may increase astral microtubule contacts with the cortex in smaller cells, potentially disrupting contractility or the coordination of chromosome segregation and cytokinesis (Rankin and Wordeman, 2010). Along with higher microtubule turnover and the uncoupling of astral and spindle microtubule growth rates, these changes may impair spindle assembly and chromosome capture (Wollman et al., 2005). Together, these effects could underlie the reduced embryonic viability in EB strains. Further analysis of astral and interphase microtubules in EB and tubulin strains will clarify how different labeling strategies influence microtubule function.

### Toward a molecular basis for the differences in microtubules dynamics between tubulin and EB strains

Microtubule growth rates are consistently lower in tubulin-tagged strains and higher in EB-tagged strains, raising the key question of whether tubulin tagging dampens growth or EB tagging enhances it. Although N-terminal tagging of tubulin could, in principle, interfere with dimer–dimer and protofilament interactions and thereby limit assembly kinetics (Honda et al., 2017), our data and previous work argue against tubulin tagging being the primary limiting factor. Introducing tagged tubulin into EB-tagged strains resulted in only a modest reduction in growth rates, with dual-labeled strains remaining much closer to EB-only than to tubulin-only conditions. Moreover, upon depletion of the microtubule polymerase ZYG-9^XMAP215^ in *C. elegans* one-cell embryos, growth rates converged to a similar basal value in both tubulin-and EB-tagged strains, regardless of the presence of tagged tubulin (Figure 4A; (Srayko et al., 2005)), indicating that tubulin tagging does not limit basal assembly. Consistently, microtubule growth rates exceeding 0.8 µm/s have been reported in oocytes expressing only tagged tubulin (Gigant et al., 2017), and spindle length and assembly timing are indistinguishable between tubulin-tagged and wild-type embryos, in contrast to EB-tagged strains, which exhibit shorter and faster-assembling spindles. Prior *in vitro* and *in vivo* cellular studies comparing labeled tubulin with label-free imaging found no differences in depolymerization rates (Luchniak et al., 2023; Cassimeris et al., 1988), suggesting again that tubulin tagging is unlikely to explain the observed differences. Finally, identical growth rates measured in *P. mammillata* embryos using either genetically encoded Venus-tubulin or microinjected chemically labeled tubulin further argue against a strong inhibitory effect of tubulin tagging, as N-terminal genetic tags and randomly distributed chemical labels would be unlikely to perturb microtubule assembly in the same manner. Together, these observations support the view that EB tagging, rather than tubulin tagging, is the primary driver of the altered microtubule growth dynamics. A recently developed split-GFP tagging of low-conservation tubulin regions minimizes functional interference and should, in principle, enable a definitive assessment of which tubulin-tagging approach is least disruptive to microtubule function (Xu et al., 2024). However, the use of an EBP-2^EB1^–mNeonGreen background in that study precludes isolating the intrinsic contribution of tubulin tagging alone.

Furthermore, tubulin-and EB-tagged strains also exhibit distinct microtubule catastrophe frequencies and depolymerization rates. The increased catastrophe frequency measured in the EB strain, which reduces average microtubule length, may explain the shorter spindles observed in this condition, despite the elevated growth rate that would otherwise predict longer spindles (Lacroix et al., 2018). Alternatively, reduced mitotic duration could contribute to the decrease in spindle length. In embryonic cleavages of most species, spindle length increases continuously from NEBD to anaphase onset (Mitchison et al., 2015), including in *C. elegans* embryos (Labbe et al., 2004; Lacroix et al., 2018). The shorter mitotic duration in the EB strain may therefore leave insufficient time for spindle assembly to reach normal length. Whether this reduction reflects altered microtubule dynamics or an effect of the tag itself remains to be investigated.

Fluorescent tags on EB proteins might locally alter protein interactions (Dörner et al., 2026). EB tagging could indeed indirectly perturb microtubule dynamics by disrupting recruitment of other plus-end tracking proteins (+TIPs), such as CLIP-170. Consistent with this view, and with our finding that CLIP-1^CLIP-170^ depletion affects tubulin strains but not EB strains (Figure 7), N-or C-terminal tagging of human EB1 abolishes CLIP-170 recruitment to microtubule plus ends (Skube et al., 2010).

Overall, our study highlights the need for caution when comparing microtubule data obtained in different contexts, particularly when distinct experimental approaches are used. This is especially important when assembling parameters from separate studies for *in silico* or mathematical modeling. Given the substantial intrinsic variability of microtubule dynamics across species and cell types (Lacroix et al., 2014), our results further suggest that the methodology used to track microtubules can itself contribute significantly to this variability.

Tubulin and EB strains highlight the plasticity of microtubule dynamics and the robustness of spindle assembly during embryonic development.

Under normal growth conditions, in absence of SAC perturbation and at optimal temperature, tubulin and EB strains exhibit similar phenotypical traits and are able to assemble functional mitotic spindle despite their highly divergent microtubule turnover. Our study thus revealed the remarkable robustness of spindle size and spindle assembly to dramatic changes in dynamics. The fact that the mitotic spindle can assemble similarly in space and in time with dynamics of individual microtubules varying almost by a factor 2 (especially growth rate and catastrophe frequency) highlights the noteworthy capacity of cells to buffer noise and variability in cellular processes and reveals the adaptability of mitotic spindle assembly. Our work provides evidence highlighting the adaptability of spindle assembly during embryonic development. Observing this adaptability in both lab-adapted *C. elegans* and wild-caught *P. mammillata* suggests that mechanisms buffering microtubule variability are evolutionarily conserved and critical for animal development, viability, and adaptation across diverse ecological niches. Evaluating microtubule dynamics relative to thermal ranges in other nematodes and ectotherms could uncover how such adaptability shapes survival, fitness, and evolutionary resilience under environmental perturbations.

## Acknowledgments

This work was supported by the Agence Nationale de la Recherche MTDiSco, (ANR-20-CE13-0033) and Ligue Nationale Contre le Cancer (Comité Département 66/ LACROIX 2022) to B.L. This work was supported by CNRS and University Paris Cité, and by grants from the European Research Council ERC-CoG ChromoSOMe 819179 and Fondation pour la Recherche Médicale FRM to J.D. Experiments on marine embryos were made thanks to an EMBRC-Fr Découverte grant in 2019 attributed to B.L. in collaboration with S.C. and J.C. We acknowledge the imaging facility MRI, member of the national infrastructure France-BioImaging (https://ror.org/01y7vt929) supported by the French National Research Agency (ANR-24-INBS-0005 FBI BIOGEN). We thank the Villefranche Service Aquariologie of CRB (IMEV-FR3761) and the Imaging Facility (PIM, a member of the MICA microscopy platform) which are supported by the EMBRC-France infrastructure network (Grant ANR-10-INBS-02). We thank the labs of Julie Arhinger, Asako Sugimoto, Bruce Bowerman, Jessica Feldman and Sander van den Heuvel for providing EB strains generated by CRISPR/Cas9 gene editing. Some worm strains were provided by the CGC, which is funded by NIH Office of Research Infrastructure Programs (P40 OD010440). We thank CRBM, Institut Jacques Monod and IMEV-LBDV lab members for their technical help and fruitful discussions.

## Experimental Procedures

### Worm culture

Worm strains were obtained from the CGC https://cgc.umn.edu or gratefully provided by different labs as mentioned in Table S1. Worm strains unless specified on the figures for the different experiments were maintained at 23°C using standard procedures (Brenner, 1974). Most of the strain used in this manuscript were outcrossed twice with wildtype strain as mentioned (“xN2×2”) in Table S1 in order to minimize the influence of different genetic backgrounds on observed phenotypes. For depletion and embryonic lethality experiments, worms were synchronized using the alkaline bleach method (1.2% NaOCl, 250 mM KOH in water (Stiernagle, 2006). Embryos collected from the alkaline bleach were allowed to hatch overnight at 16°C in M9 buffer (22 mM KH2PO4, 42 mM Na2HPO4, 86 mM NaCl) under gentle rotation. This overnight incubation allowed all intact embryo to hatch and give rise to L1 arrested worms.

### Embryonic lethality assays

Synchronized L1 larvae were allowed to grow on feeding plate at the indicated temperature until adulthood. Since development timing is dependent on temperature, the onset of adulthood for this experiment was considered when the first laid embryos start to hatch on the plate. Since worms are synchronized the presence of hatched embryo indicate that almost all worms on the plate had reached the adult stage. An individual adult was then transferred onto a new plate and allowed to lay eggs for 24h before being removed from the plate. The day after, the number of larvae (hatched eggs) and the number of embryo (dead progeny) was counted manually. The percentage of lethality correspond to 100 x (dead progeny) / (total progeny). 5 biological and 3 experimental replicates were done for this experiment.

### Protein depletion in *C. elegans*

Protein depletion was performed as described previously (Kamath et al., 2001) by feeding worms with HT115 bacterial strain containing the L4440 vector allowing for for IPTG mediated induction of dsRNA expression. Individual bacterial clones from the Arhinger’s library were kindly provided by Jean-Claude Labbé (IRIC, Université de Montréal) or Lionel Pintard’s lab (Institut Jacques Monod, Paris) and targets of all clones used in this study were confirmed by sequencing. A single colony of transformed HT115 bacteria was incubated for 7-8h in 2 mL of LB medium supplemented with 50 µg/mL carbenicillin. The bacterial culture was then seeded onto NGM plates containing 0.2 µM IPTG (Isopropyl *β*-D-thiogalactopyranoside) and 50 µg/mL of carbenicillin and incubated overnight at room temperature. Between 30 and 50 L1 worms were placed on a NGM plate seeded with the culture of HT115 bacteria expressing dsRNA and allowed to grow in the dark at the indicated temperature.

### Worm mounting and imaging conditions

Gravid worms were dissected in egg salt buffer (ESB, 25 mM HEPES pH 7.3, 118 mM NaCl, 48 mM KCl, 2 mM CaCl2, 2 mM MgCl2) by cutting each worm open using a 26-gauge syringe needle. Embryos were then transferred on a 3% agarose pad and mounted between a glass slide and a coverslip. The montage was then sealed with Valap (vaselin:lanolin:parrafin wax, 1:1:1 weight). The imaging chamber was filled with ESB buffer to avoid drying. Nomarski (DIC) images were acquired on a Zeiss Axioimager Z2 microscope with a 63 Plan Apochromat 1.4 NA oil immersion objective. Kölher alignment was performed prior to every imaging session. The first zygotic division was followed by timelapse during 10-15 min at 23°C with 5 s interval. Movies to track microtubule dynamics were acquired on a spinning disc confocal microscope. To minimize toxicity, we used spinning-disk (CSU-X1, Yokogawa) confocal microscope (Roper Scientific) coupled to a CoolSnap HQ2 CCD camera (Photometrics).

Microscope and acquisitions were controlled by MetaMorph 7 software (Molecular Devices). Movies recorded to track microtubule dynamics correspond to a single focal plane imaged at 2 frame per second interval using a Nikon CFI APO LBDA S 60x/NA1.4 oil objective with no camera binning. Observation of chromosome segregation effect were done by imaging developing embryos under a spinning disk confocal Yokogawa W1 on an Olympus inverted microscope coupled to a sCMOS Fusion BT Hamamatsu camera. Individual embryos were imaged for a maximal duration of 10 min to minimize photo damage. A z-stack of 15 to 20 µm with a z-section of 1 µm was made every 10 second using a 60× UPLSAPO 1.3 NA DT 0.3 mm silicone objective.

### Scoring frequency of lagging chromosomes

3D time-lapse of *C. elegans* embryo imaged in the channel for fluorescent histone (mCherry-Histone in different background) were visualized in ImageJ. The number of chromosomal defects: misaligned chromosome in metaphase and lagging chromosome in anaphase were scored manually at each stage of the development. Developmental stage was estimated by counting the number of blastomere on the entire z-stack. Individual embryos were imaged only during a single round of division to avoid accumulation of defects that would be due to laser illumination and imaging conditions. As the number of cells is doubled at each stage, the frequency of defect was estimated per observed dividing cells, not per embryo. At least 12 embryos were imaged and analyzed for each stage.

### Tubulin preparation and fluorescent labeling

Tubulin was obtained from pig brains following cycles of polymerization and depolymerization (Castoldi and Popov, 2003). Tubulin was then labeled with NHS-ester-ATTO 488 or NHS-ester-ATTO 565 (ATTO-TEC) and further purified through two polymerization/depolymerization cycles (Hyman, 1991) to ensure that only tubulin dimer able to assemble into microtubules are purified.

### Ascidian egg fertilization and embryo culture

Adult *Phallusia mammillata* were collected at Sète (Etang de Tau, Mediterranean coast, France) or Roscoff (Brittany, France) and maintained for several months in aquaria at the Centre de Ressources Biologiques (CRB) of the Institut de la Mer à Villefranche (IMEV). Following dissection of the adult hermaphrodite, gametes were collected with pipets from the oviduct and spermiduct. Eggs were de-chorionated by treatment with 0.1% trypsin (Sigma-Aldrich, T9201) in micro-filtered seawater (MFSW) buffered with 5 mM TAPS pH 8.2 ([tris (hydroxymethyl) methylamino] propanesulfonic acid) for 1-2 hours. Because dechorionated eggs adhere to glass and plastic, all dishes and pipettes are coated with agarose or “GF” (a mixture of 0.1% gelatin and 0.1% formaldehyde) (Sardet et al., 2011; McDougall et al., 2015). To fertilize, sperm was activated by dilution into pH 9.5 sea water for 10 minutes then added to eggs at a ratio of ∼1: 100. Once eggs changed shape which indicates fertilization, they were transferred to fresh MFSW to prevent polyspermy. Embryos were then cultured in GF-coated dishes at 18°C until the desired stage.

### Microinjection of *Phallusia mammillata* eggs

Dechorionated eggs were placed in horizontal glass injection chambers filled with MFSW (Sardet et al., 2011; Chenevert et al., 2024). Injection needles were made from thin wall glass capillaries without filament, inner diameter 0.78 mm outer diameter 1 mm (GC100T-10, Harvard Apparatus) using a Narishige PN-30 horizontal puller. Needles are placed in a HI-7 needle holder which receives pressure from a Narishige IM300 injection box attached by oil-filled tubing to a Mecafer air compressor. The needle is controlled by a three-axis hydraulic micromanipulator (Narishige MMO-203) mounted on a Nikon DiaPhot inverted microscope fitted with 10X and 20X objectives.

Needles were front-filled from injection tubes prepared from glass capillaries (GC100T-10, Harvard Apparatus) containing 1 µl RNA flanked by 1 µl layers of mineral oil (Sigma). The concentration of RNA in the needle was 1-5 μg/μl and the amount injected was 2-5% egg volume as estimated by the small clearing in the cytoplasm. RNAs were synthesized from pRN3-derived plasmids harboring GFP sequence fused to the C-terminus of human EB3 (GenBank accession number AY893969.1 (Rosfelter et al., 2024) or Venus sequence fused to the N-terminus of tubulin beta. EB3-GFP and venus-tubulin RNAs were transcribed using T3 RNA polymerase and 5’ capped with the mMessage mMachine kit (Ambion). Injected eggs were left in MFSW at 18°C for several hours or overnight before fertilization and imaging.

### Imaging of Phallusia mammillata embryos

Imaging was performed with a Leica TCS SP8 inverted confocal microscope fitted with a 40×/1.1NA water immersion objective and 488 and 552 nm lasers and Leica application suite acquisition software (LASX). For mounting of live samples, embryos were placed in a drop of MFSW in GF-coated glass bottom dishes (Cellvis) secured with a coverslip as described (Chenevert et al., 2024). A focal plane was imaged for the relevant cells was acquired in both fluorescent and brightfield channels with a time interval of 862 ms (1.16 image per second). All live imaging experiments of *Phallusia mammillata* embryos were performed at 18–19 °C.

### Image processing and analysis of microtubule dynamics

Timelapse movies were converted into non-compressed.avi files and processed in ImageJ software (https://imagej.net). Microtubule dynamics were extracted as described in Chenevert et al. (Chenevert et al., 2024). Kymographs were generated using an in ImageJ macro available at https://github.com/benlacroix/ImageJ_Macros allowing for semi-automated generation of kymographs and based on the ImageJ plugin’Multiple kymograph’ (http://www.embl.de/eamnet/html/kymograph.html).

Dynamic parameters were extracted from kymographs using trigonometry calculation available here on: https://github.com/benlacroix/MTDynamX_Data_Xtraction. Scatter plots, histograms and statistical analyses were generated and performed using Prism (GraphPad Software). Means were compared using the unpaired Welch t test. Diamond graphs were drawn using a custom MATLAB-based software (MathWorks) on which rates (growth and shrinkage) and frequencies (catastrophe and rescue) were jointly normalized as they are expressed in the same.

## CONTACT FOR REAGENT AND RESOURCE SHARING

Benjamin Lacroix, Centre de Recherche en Biologie Cellulaire de Montpellier, France

