## Supplemental Data for "The extraordinary robustness of mitotic spindle assembly to microtubule dynamics plasticity revealed by end-binding proteins and tubulin tagging"

#### Figure S1

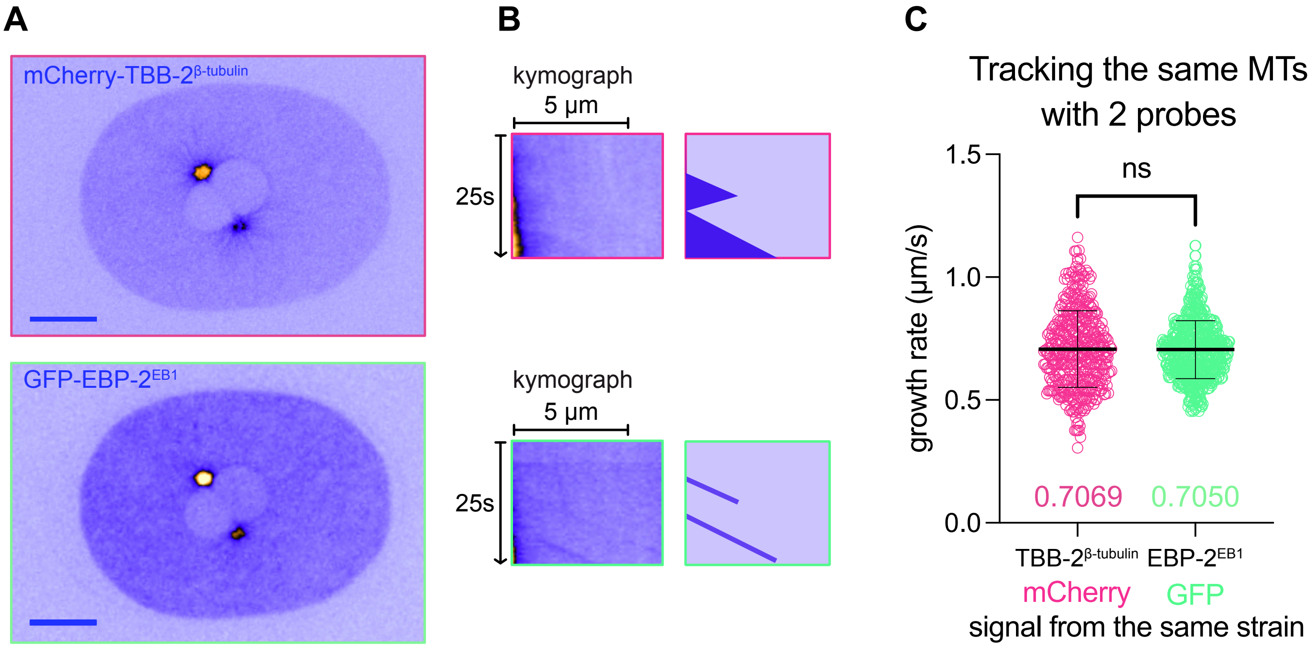

A
mCherry-TBB-2^β-tubulin
GFP-EBP-2^EB1
B
kymograph
5 µm
25s
kymograph
5 µm
25s
C
Tracking the same MTs with 2 probes
1.5
1.0
0.5
0.0
growth rate (µm/s)
ns
0.7069
0.7050
TBB-2^β-tubulin
EBP-2^EB1
mCherry
GFP
signal from the same strain

**Figure S1. Related to Figure 1. Fluorescent EB and tubulin decorate the same microtubules.**

(A) Snapshots extracted from a 2-color time lapse video of an embryo expressing both mCherry-tubulin (top panel, red channel) and GFP-EBP-$2^{EB1}$(bottom panel, green channel). Scale bars = 10 µm. (B) Kymographs extracted from a single microtubule trajectory highlight the same dynamic events visualized by both markers. Tubulin labeling allows the visualization of polymerization and depolymerization events while EBP-$2^{EB1}$ marker only highlights growth excursions. (C) Microtubule growth rate measured independently from the same movies using either tubulin marker or EB marker. The average growth rate was identical between the two independent measurements, ns: no significant difference using Welch's t test comparison.

#### Figure S2

**A**

$$J = \frac{v_g \cdot f_r - v_s \cdot f_c}{f_c + f_r}$$

**B**

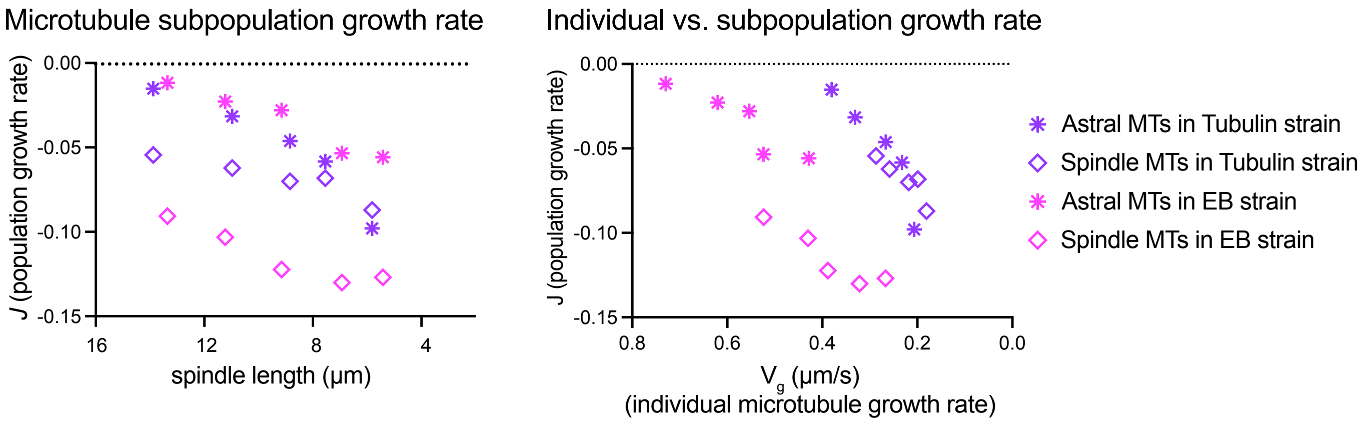


**C**

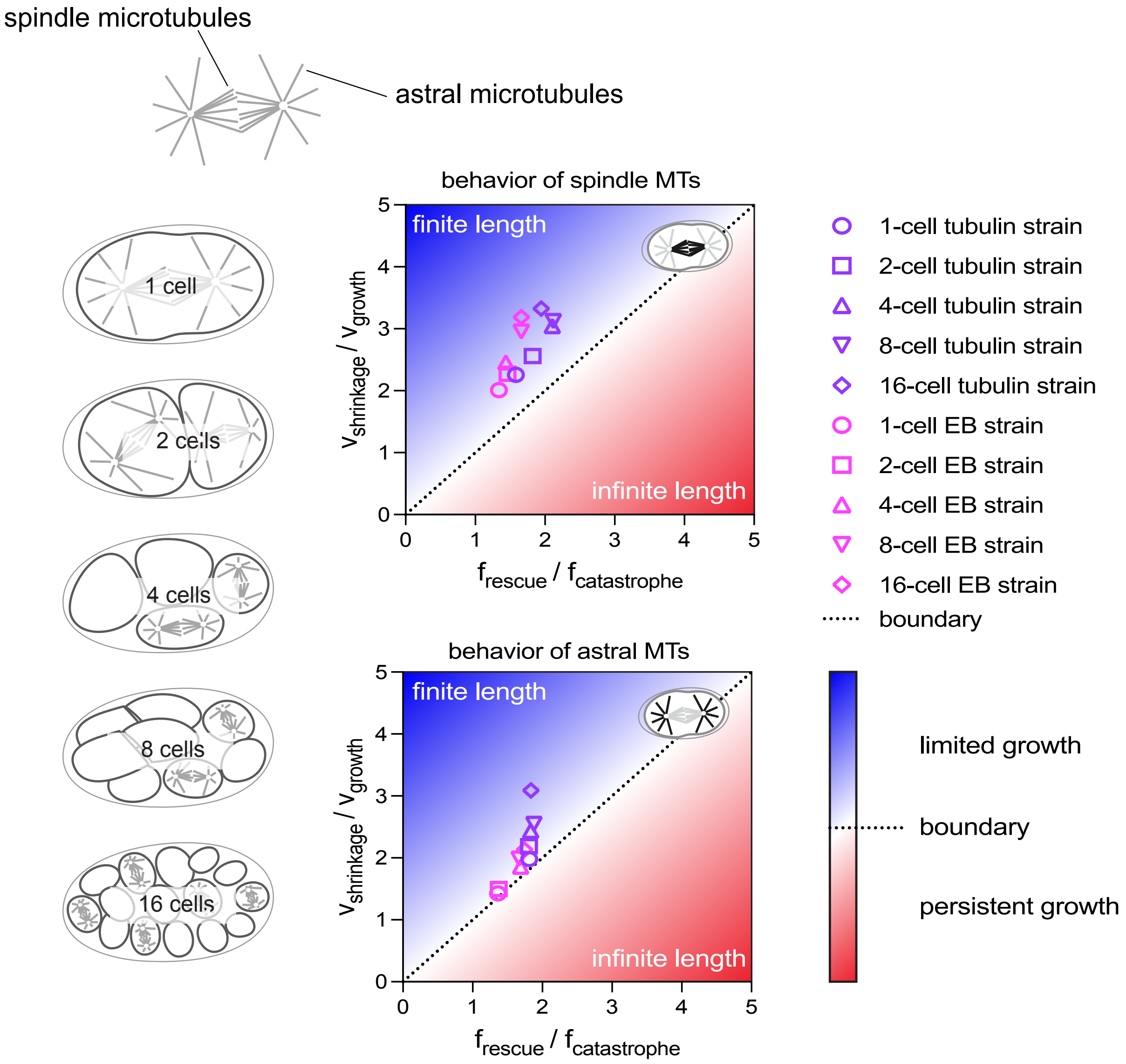

**Figure S2. Related to Figure 3 and to the Discussion. Microtubule population growth rate and phase diagrams of microtubule dynamics during embryonic cleavages in tubulin strains and EB strains.**

(A) The equation from Verde et al. (Verde et al., 1992) used to estimate the growth rate of a microtubule population "$J$" based on the four parameters of microtubule dynamics. (B) The parameter *J* was plotted against spindle length (left) or individual microtubule growth rate (right) for astral and spindle microtubule and from 1-cell to 16-cell stage embryos. If the ratio $f_r/f_c$ is greater than the ratio $v_s/v_g$, $J$ becomes positive and microtubules tend to growth infinitely (unbounded). The dashed line at *J*=0 represent the boundary. (B) Phase diagrams to visualize microtubule behavior relative to the boundary between infinite length and steady-state length. Parameters from Figure 3 were used to report $f_r/f_c$ and $v_s/v_g$ for spindle (top diagram) and astral (bottom diagram) microtubules from 1-cell to 16-cell stage embryos (drawings on left). In both tubulin strains and EB strains, spindle and astral microtubule dynamics remain within the bounded regime during the mitotic phase. However, we noticed that astral microtubules in EB strain are very close to the boundary. These dynamic properties might result in increased number of astral microtubules contacting the cortex in small cells that could lead to defects in contractility or coordination between mitosis and cytokinesis (Rankin and Wordeman, 2010).

### Figure S3

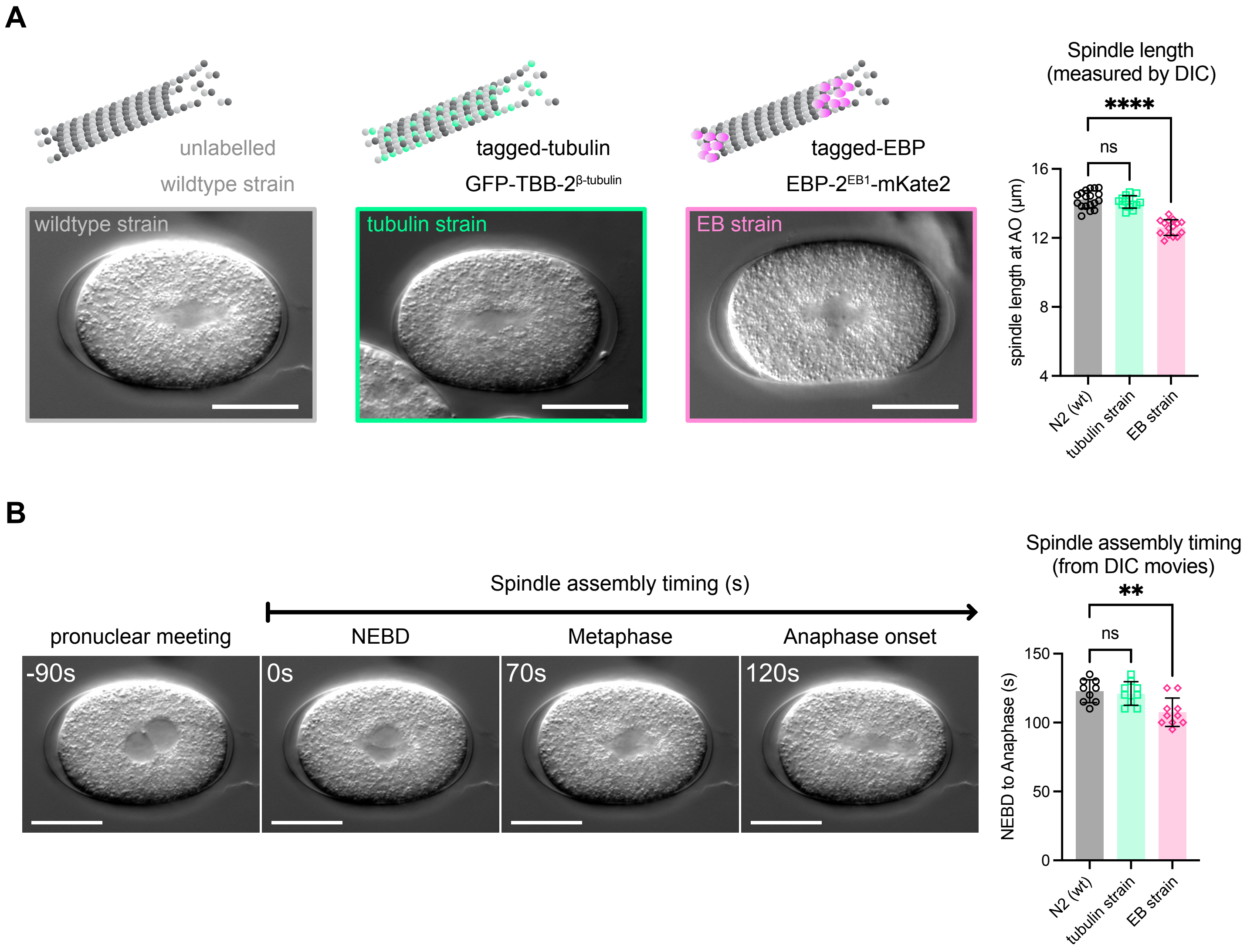

A
unlabelled
wildtype strain
tagged-tubulin
GFP-TBB-2β-tubulin
tagged-EBP
EBP-2EB1-mKate2
wildtype strain
tubulin strain
EB strain
Spindle length
(measured by DIC)
****
ns
spindle length at AO (µm)
16
12
8
4
N2 (wt)
tubulin strain
EB strain
B
Spindle assembly timing (s)
pronuclear meeting
NEBD
Metaphase
Anaphase onset
-90s
0s
70s
120s
Spindle assembly timing
(from DIC movies)
**
ns
NEBD to Anaphase (s)
150
100
50
0
N2 (wt)
tubulin strain
EB strain

**Figure S3. Related to Figure 5. Spindle size and assembly duration is influenced by microtubule labelling method.**

(A) Differential interference contrast (DIC) images of live *C. elegans* 1-cell embryos from a wildtype N2 strain, a tubulin strain (GFP-TBB-$2^{tubulin}$) and an EB strain (EBP-$2^{EB1}$-mKate2). Scale bar = 20 µm. Spindle length was measured at anaphase onset from at least 14 embryos. Statistical differences were evaluated by a Dunnett's multicomparison test. ****: P value <0.0001, ns: no significant difference. Colored bars represent mean and black whiskers indicate standard deviation. (B) Spindle assembly duration evaluated from DIC video microscopy images. The bar graph represents the average duration from at least 9 different embryos per condition. Colored bars represent mean and black whiskers indicate standard deviation. **: P value <0.01, ns: no significant difference using Dunnett's multicomparison test.

### Table S1

| Nomenclature used in the manuscript for fluorescent markers | Type of strain (for this study) | Genotype | Worm strain | Source | Comments |
|---|---|---|---|---|---|
| **N2 (wt)** | wildtype | wildtype | N2 (Bristol) | CGC https://cgc.umn.edu/ | Commonly used wild type strain |
| **GFP-TBA-2$^{\alpha\text{-tubulin}}$** | tubulin strain | *ijmSi63* [pJD520; mosII_5'mex-5_GFP::*tba-2*; mCherry::*his-11*; cb-unc-119(+)] II; *unc-119(ed3)* III? | JDU233 | Julien Dumont's lab | tubulin expressed under germline specific promoter (mex-5) |
| **mCherry-TBB-2$^{\beta\text{-tubulin}}$** | tubulin strain | [pJA138; 5'pie-1::mCherry::tbb-2::3'pie-1] | JA1559* | Julie Ahringer's lab | tubulin expressed under germline specific promoter (pie-1) |
| **EBP-1$^{EB1}$-GFP** | EB strain | unc119(ed3) III; tjls8 [pie-1::GFP::ebp-1, unc-119(+)] | SA141 | Fumio Motegi, Asako Sugimoto's lab | EBP expressed under germline specific promoter (pie-1) |
| **mCherry-TBB-2$^{\beta\text{-tubulin}}$ ; EBP-1$^{EB1}$-GFP** | EB strain | *unc119(ed3)* III; *tjls8* [pie-1::GFP::*ebp-1, unc-119*(+)]; [pJA138; 5'pie-1::mCherry::tbb-2::3'pie-1] | JDU400* | Julien Dumont's lab Made by crossing SA141 males with JA1559 | EBP expressed under germline specific promoter (pie-1) |
| **EBP-2$^{EB1}$-GFP(#1)** | EB strain | ebp-2(wow47[ebp-2::GFP])II | JDU695* | Julien Dumont's lab. Made by outcrossing JLF273 from Jessica Feldman's lab with N2 | EBP tagged by CRISPR/Cas9 gene editing |
| **mCherry-TBB-2$^{\beta\text{-tubulin}}$ ; EBP-2$^{EB1}$-GFP(#1)** | EB strain | [pJA138; 5'pie-1::mCherry::tbb-2::3'pie-1] I ?? ; ebp-2(wow47[ebp-2::GFP])II; | JDU692* | Julien Dumont's lab. Made by crossing JLF273 from Jessica Feldman's lab with JA1559 | EBP tagged by CRISPR/Cas9 gene editing |
| **GFP-TBB-2$^{\beta\text{-tubulin}}$ ; EBP-2$^{EB1}$-mKate2** | EB strain | *ebp-2[or1954(ebp-2::mKate2)]II, ruls57 [pAZ147: pie-1p::GFP::tubulin; unc-119 (+)]* | EU3068* | Bruce Alan Bowerman'lab | EBP tagged by CRISPR/Cas9 gene editing |
| **GFP-TBB-2$^{\beta\text{-tubulin}}$** | tubulin strain | *ruls57 [pAZ147: pie-1p::GFP::tubulin; unc-119 (+)]* | JDU598* | EU3068 outcrossed with N2 to remove EB marker | tubulin expressed under germline specific promoter (pie-1) |
| **EBP-2$^{EB1}$-mKate2** | EB strain | *ebp-2*[*or1954*(*ebp-2*::mKate2*)*]II. | JDU597* | EU3068 outcrossed with N2 to remove tubulin marker | EBP tagged by CRISPR/Cas9 gene editing |
| **EBP-2$^{EB1}$-GFP(#2)** | EB strain | ebp-2[he??(ebp-2::gfp)]II | SV1937* | Sander Jean Louis van den Heuvel's lab | EBP tagged by CRISPR/Cas9 gene editing |
| **« wt » (with histone marker)** | - | unc-119(ed3) III; itls37[pAA64; pie-1/ mCherry::his-58; unc-119(+)] IV | OD56* | Julien Dumont's lab | - |
| **« EB strain » (with histone marker)** | EB strain | ebp-2(wow47[ebp-2::GFP])II; unc-119(+)]III; ltls37[pAA64; pie-1/mCHERRY::his-58; unc-119(+)]IV. | JDU696* | Julien Dumont's lab | EBP tagged by CRISPR/Cas9 gene editing |
| **« tubulin strain » (with histone marker)** | tubulin strain | ruls57 [pAZ147: pie-1promoter::tubulin::GFP; unc-119 (+)];unc-119(ed3) III; itls37[pAA64; pie-1/mCherry::his-58; unc-119(+)] IV | Unamed* | This study | tubulin expressed under germline specific promoter (pie-1) |

* these strains were outcrossed x2 with N2 (wt) worms in our hands to reduce the effect of divergent genetic background on our results.
